# Single-dose rapamycin increases brain glucose metabolism but reduces synaptic density in Long- Evans rats

**DOI:** 10.1101/2025.03.11.642549

**Authors:** Frederik Gudmundsen, Zhen Li, Christina Baun, Jonas E. Svensson, Niels Langkjær, Mikael Palner, Pontus Plavén-Sigray

## Abstract

Rapamycin, an inhibitor of the mechanistic target of rapamycin (mTOR), has shown promise as a neuroprotective compound in preclinical studies. Reduced brain glucose metabolism and loss of synaptic density are key features of Alzheimer’s disease that can be measured *in vivo* using positron emission tomography (PET) imaging, allowing for assessment of treatment effects on brain function. Here, we used PET to investigate the acute effects of a single-dose of rapamycin on glucose metabolism and synaptic density in Long-Evans rats.

In a repeated measures design, we quantified changes in brain glucose metabolism using [^18^F]FDG PET (n=13) at baseline, one day, and one week after intraperitoneal administration of rapamycin (8 mg/kg). In a separate cohort (n=6), we measured synaptic density using [^18^F]SynVesT-1 PET at baseline and one day after rapamycin administration. Regional standardized uptake values (SUV) were calculated for [^18^F]FDG while total distribution volumes were estimated for [^18^F]SynVesT-1 using image-derived input functions of the heart.

Rapamycin induced significant increases in [^18^F]FDG SUV across multiple brain regions one day after administration, an effect that persisted at one-week follow-up. In contrast, [^18^F]SynVesT-1 binding showed significant decreases throughout the brain at 24 hours post-administration, indicating reduced synaptic density. These opposing effects on glucose metabolism and synaptic density point to multifaceted actions of rapamycin in the brain, possibly reflecting improved metabolic function occurring simultaneously with acute synaptic loss.

These results show that [^18^F]FDG and synaptic density PET imaging could serve as useful biomarkers in human clinical trials evaluating rapamycin’s mechanistic and therapeutic effects in neurodegenerative disorders.

## INTRODUCTION

For decades, amyloid-targeted therapies have been the dominant approach in Alzheimer’s disease (AD) drug development. However, clinical trials targeting amyloid have shown mixed results[1], with only recent approvals of antibody treatments that remain limited by cost, eligibility and safety concerns [2– 4]. This has highlighted the need to explore alternative therapeutic strategies, particularly those targeting fundamental aging-related mechanisms that contribute to neurodegeneration [5,6].

Rapamycin, an inhibitor of the mechanistic target of rapamycin (mTOR), shows potential as a neuroprotective treatment for age-related conditions, such as AD [7]. In preclinical studies, rapamycin extends lifespan across multiple species and delays the onset of various age-related pathologies [8]. Notably, treatment with rapamycin or analogous mTOR inhibitors in mouse models of AD has been shown to reduce amyloid-β deposition, decrease pathogenic tau phosphorylation, improve cerebral blood flow, and enhance cognitive function [6,7]. These effects have been observed both when treatment is initiated before symptom onset and after pathology is established suggesting potential use in both prevention and treatment.

Brain glucose metabolism, measured by [^18^F]FDG positron emission tomography (PET), is a well-established biomarker of neuronal function and has proven valuable for early detection and monitoring of AD progression [9,10]. Changes in [^18^F]FDG PET uptake have previously been used as endpoint in early-phase clinical trials of candidate AD medications [11]. More recently, PET imaging of synaptic vesicle glycoprotein 2A (SV2A) [12–14] has enabled *in vivo* quantification of synaptic density, providing new opportunities to track synapse loss [15,16], which is a key feature of neurodegeneration [17,18] that correlates strongly with cognitive decline [19]. By combined assessment of these markers, we can get a better understanding for how brain metabolism and synaptic integrity is affected by potential neuroprotective drugs like rapamycin.

The present study investigated the effects of single-dose rapamycin administration on brain glucose metabolism and synaptic density in Long-Evans rats using [^18^F]FDG and [^18^F]SynVesT-1 PET imaging, respectively. This approach allows assessment of rapamycin’s acute effects on both metabolic function and synaptic integrity, and can serve to inform the design of upcoming clinical studies evaluating rapamycin’s effects in patients with neurological disorders, such as AD, where imaging markers are to be used as trial endpoints [20].

## METHODS

### Animals

Young-adult Long Evans rats (n=24) were purchased from Janvier labs (Le Genest-Saint-Isle, France) and maintained under a 12h light/dark cycle (6 a.m. to 6 p.m.) with unrestricted access to food (Altromin 1324, Brogaarden) and water. To achieve optimal acclimation and habituation, rats arrived at the animal facility at least one week prior to experiments. Rats were housed in groups of 4 in ventilated cages with controlled temperature (22-24 °C) and relative humidity (55 ± 10%). Environmental enrichment was provided, including red-colored shelters, wooden sticks, and bedding (Classic Aspen Bedding 2-5mm, Scanbur). All rats were moved to the PET imaging facility 24h prior to experiments for acclimation to the scan room environment. In the PET facility, animals were housed in conventional cages with dust-free bedding (Pure-O’Cell, Scanbur) within a Scantainer (EHRET) to maintain consistent temperature and humidity (same as above). All procedures were conducted in accordance with the FELASA guidelines and with approval from the Danish Animal Experiments Inspectorate (license number: 2021-15-0201-00909).

### Production of PET radioligands

Two different PET radioligands were used to assess brain glucose metabolism and synaptic density, [^18^F]FDG and [^18^F]SynVesT-1, respectively. [^18^F]FDG was produced at the Department of Clinical Physiology and Nuclear Medicine, Odense University Hospital, as part of their daily clinical productions with a purity exceeding 99% and a molar activity of 33 GBq/μmol.

[^18^F]SynVesT-1 was synthesized for this study via direct radiofluorination of its trimethylstannyl precursor (Pharmasynth, Tartu, Estonia) using a Fastlab Gen 2 with FASTlab HPLC+ (GE Healthcare, Uppsala, Sweden). No-carrier-added [^18^F]fluoride was produced by a PETtrace cyclotron (GE Healthcare) through proton irradiation of [^18^O]H_2_O (Rotem Medical) and isolated on a pre-conditioned QMA cartridge (Waters Sep-Pak). The trapped [^18^F]fluoride was dried using nitrogen flow followed by dry DMA wash to remove residual water. The fluoride was eluted with Cu(OTf)_2_ (0.0645 M in DMA, 700 μL) into a reaction vial containing the precursor (2 mg), pyridine (50 μL), and DMA (450 μL). The reaction proceeded at 110°C for 20 minutes. The diluted crude mixture underwent semi-preparative HPLC purification (Phenomenex Luna C18 column) using 20% ethanol in water with phosphoric acid. The product eluted at ∼13 min, yielding 2540 MBq of [^18^F]SynVesT-1 ready for immediate use. Analytical HPLC confirmed >99% radiochemical purity with 11.7 GBq/μmol molar activity

### [^18^F]FDG imaging

To measure the effect of rapamycin on brain glucose metabolism, a series of [^18^F]FDG-PET recordings were obtained in 16 young-adult Long Evans rats (three excluded, see below) in a single-arm three-scan repeated measure study: baseline, one day post-treatment, and one-week post-treatment. Animals received saline (400 μl, intraperitoneal; I.P.) before baseline and one-week post scans, and rapamycin (8 mg/kg in 5% ETOH in PEG:Tween:Water solution, I.P.) 24h prior to the one-day post scan. This dose was chosen to correspond to dosing regimens showing beneficial neuro or geroprotective effects in previous rodent studies [21,22].

The scanning procedures followed previous protocols that has been described in detail elsewhere [23]. In brief, for each scan occasion, rats were fasted overnight prior to the [^18^F]FDG-PET recording to deplete glucose levels. Prior to each scan, rats were weighed and blood glucose was measured (Baseline mean = 6.70 ± 1.03 mmol/L; one-day post mean = 6.77 ± 0.82 mmol/L; one-week post mean = 6.70 ± 0.57 mmol/L, all *p* > 0.78). Subsequently, rats were anesthetized with isoflurane (induction 4%, 1-2% maintenance) and [^18^F]FDG (8.6 ± 1.7 MBq) was administered through a bolus IV injection. Following [^18^F]FDG administration, animals were returned to their home cage to allow awake radioligand uptake. After 45 min, the animals were lightly anesthetized and placed in a Siemens INVEON multimodality preclinical scanner in docked mode (Siemens, Knoxville, TN, US) for 35 min dynamic recording (5 min CT followed by a 30 min list-mode emission scan). While in the scanner, rats were positioned on a heating plate to maintain a stable body temperature of 37 °C with respiration monitoring using the BioVet system (M2M imaging, Cleveland, Ohio, US).

The CT scan was performed with the following parameters: 270-degree full rotation, 4×4 binning, low magnification, and a 90 mm field of view (FOV), with exposure settings of 80 kV and 500 μA for 350 ms. The CT scans were reconstructed with the Feldkamp algorithm using a Shepp Logan filter and Hounsfield calibration, resulting in images with an effective pixel size of 111.25 μm, which was used for attenuation correction and anatomic registration. Emission data were reconstructed into a single static image using the ordered subset expectation maximization 3D (OSEM3D/SP-MAP) algorithm (2 × OSEM iterations and 18 × MAP iterations) applying decay and scatter correction in a 128 × 128 matrix of 1.4 mm pixels.

For the [^18^F]FDG experiments, two rats were excluded from the experiment due to emission reconstruction failure and one rat was excluded due to injection failure at baseline. In total, 13 rats (weight: 355.5 ± 13.8 g) were included in the final analysis.

### [^18^F]FDG analysis

After image acquisition, attenuation correction and reconstruction, [^18^F]FDG-PET images were head cropped, and the averaged brain uptake images were co-registered to the Schiffer [^18^F]FDG template for segmentation by the Schiffer volumes of interest atlas in PMOD (v. 4.2, PMOD Technologies, Switzerland). The decay corrected [^18^F]FDG uptake (kBq/cc) was converted to standardized uptake values (SUV) with respect to bodyweight and injected dose.

### [^18^F]SynVesT-1 imaging

A separate cohort of eight young-adult Long Evans rats (two excluded, see below) underwent [^18^F]SynVesT-1-PET imaging to assess rapamycin’s effect on synaptic density. In a single-arm design, two repeated scans were conducted on each animal: at baseline (24h after saline administration, 400μl, I.P.) and at follow-up (24h after rapamycin administration, 8 mg/kg in 5% ETOH in PEG:Tween:Water solution, I.P.).

Rats were anesthetized with isoflurane (induction 4%, maintenance 1-2%) and placed in the Siemens Inveon Micro PET/SPECT/CT scanner for a 5min CT attenuation scan. After insertion of an IV catheter in the tail vein, [^18^F]SynVesT-1 (17.4 ± 2.2 MBq) was administered as a bolus injection at scan start. A 90-min dynamic scan was acquired for each rat, using the same scan and reconstruction specifications as for [^18^F]FDG experiments. The 90-min emission scans were then reconstructed into 23 frames (6×10s, 8×30s, 5×300s, 4×900s).

For the [^18^F]SynVesT-1 experiments, two rats were excluded from the experiment due to synthesis failure. In total, 6 rats (weight: weight: 294.9 ± 34.9 g) were included in the final analysis.

### [^18^F]SynVesT-1 analysis

Reconstructed [^18^F]SynVesT-1 PET images were analyzed using the same pre-processing pipeline as the [^18^F]FDG-PET images. For SUV analysis, decay-corrected activity was converted with respect to bodyweight and injected dose, with regional SUV calculated as an average from 27.5 min to the end of scan. For estimation of total distribution volume (VT), time activity curves were extracted for all regions, and total blood activity was estimated through an image-derived input function (IDIF) from the heart. Regional estimates at baseline and 24h post rapamycin was quantified using a Logan plot with the extracted IDIF. Since these IDIFs consist of a mixture of venous and arterial blood and are not corrected for metabolites, we denote them as “pseudo-VT” estimates below.

### Statistics

Regional differences in SUV or pseudo-VT values between conditions were tested through two-way ANOVA (α=0.05), with adjustment for multiple comparisons using Tukey’s method for multiple comparisons. Differences in global uptake and binding in a composite mask consisting of (volume-weighted) regions were assessed using Wilcoxon non-parametric test (α=0.05).

## RESULTS

### [^18^F]FDG

For [^18^F]FDG (n=13), there was a significant increase in [^18^F]FDG SUV in all individual regions following injection of rapamycin (one-day post, all p< 0.05, see Figure 1 and Table 1), but not in a volume weighted composite-mask of all regions (p = 0.13, 9.3%). This change persisted one week following administration of the drug (Figure 1, Table 1), and was also significant in the composite mask (p = 0.04, 14.5%). Regional increase in SUV ranged from 7.5 to 11.8% for one-day post, and from 12.1-17.0%, for one-week post, with the strongest effect observed in the hippocampus medial prefrontal cortex, thalamus and somatosensory cortex, albeit with some variability between one day and one week post follow-up scans (see Table 1).

**Table 1.** Average % change (“%”) in [^18^F]FDG uptake and [^18^F]SynVesT-1 binding as proxies for glucose metabolism and synaptic density, respectively, following a single I.P. dose of rapamycin.

| Regions | [<sup>18</sup>F]FDG SUV |  |  |  |  |  |  | [<sup>18</sup>F]SynVesT-1 pseudo-V<sub>T</sub> |  |  |  |
| --- | --- | --- | --- | --- | --- | --- | --- | --- | --- | --- | --- |
|  | Baseline | Acute effects |  |  | One week effects |  |  | Baseline | Acute effects |  |  |
|  |  | One day follow-up | % | *p* | One-week follow-up | % | *p* |  | One day follow-up | % | *p* |
| NAc | 7.32 ± 1.34 | 7.78 ± 1.02 | 7.5 | <0.0001 | 8.11 ± 0.82 | 12.1 | <0.0001 | 6.24 ± 0.37 | 5.74 ± 0.50 | -7.9 | 0.0018 |
| Amyg | 5.38 ± 1.11 | 5.95 ± 0.89 | 10.6 | <0.0001 | 6.11 ± 0.53 | 13.6 | <0.0001 | 5.22 ± 0.38 | 4.80 ± 0.53 | -8.0 | 0.0159 |
| ACC | 7.66 ± 1.62 | 8.84 ± 1.41 | 10.7 | <0.0001 | 8.79 ± 1.04 | 14.8 | <0.0001 | 5.45 ± 0.26 | 5.09 ± 0.51 | -6.5 | 0.0731 |
| EntoC | 5.92 ± 1.20 | 6.49 ± 0.99 | 9.6 | <0.0001 | 6.71 ± 0.63 | 13.4 | <0.0001 | 4.95 ± 0.42 | 4.54 ± 0.50 | -8.2 | 0.0204 |
| HIP AD | 6.08 ± 1.26 | 6.77 ± 0.98 | 11.4 | <0.0001 | 7.02 ± 0.73 | 15.5 | <0.0001 | 5.29 ± 0.27 | 5.11 ± 0.49 | -3.4 | 0.9229 |
| HIP post | 5.45 ± 1.12 | 5.95 ± 0.87 | 11.8 | <0.0001 | 6.20 ± 0.70 | 13.8 | <0.0001 | 5.39 ± 0.27 | 5.02 ± 0.49 | -6.8 | 0.0537 |
| Hypotha | 5.49 ± 1.03 | 5.95 ± 0.83 | 8.4 | <0.0001 | 6.19 ± 0.59 | 12.8 | <0.0001 | 6.13 ± 0.40 | 5.64 ± 0.50 | -8.1 | 0.0017 |
| Insula | 7.04 ± 1.47 | 7.57 ± 1.21 | 7.5 | <0.0001 | 8.03 ± 0.74 | 14.0 | <0.0001 | 5.71 ± 0.29 | 5.22 ± 0.44 | -8.6 | 0.0020 |
| mPFC | 8.04 ± 1.61 | 8.95 ± 1.26 | 11.4 | <0.0001 | 9.08 ± 0.90 | 13.0 | <0.0001 | 6.17 ± 0.29 | 5.74 ± 0.47 | -7.0 | 0.0099 |
| Midbrain | 6.58 ± 1.18 | 7.17 ± 0.94 | 9.0 | <0.0001 | 7.59 ± 0.74 | 15.4 | <0.0001 | 6.19 ± 0.33 | 5.67 ± 0.50 | -8.4 | 0.0009 |
| Olfact. | 6.96 ± 1.38 | 7.53 ± 1.13 | 8.2 | <0.0001 | 7.82 ± 0.73 | 12.4 | <0.0001 | 5.36 ± 0.34 | 4.91 ± 0.50 | -8.3 | 0.0071 |
| OFC | 7.89 ± 1.57 | 8.57 ± 1.31 | 8.7 | <0.0001 | 8.90 ± 0.96 | 12.8 | <0.0001 | 5.07 ± 0.15 | 4.69 ± 0.44 | -7.6 | 0.0344 |
| SSC | 6.77 ± 1.31 | 7.38 ± 1.21 | 9.1 | <0.0001 | 7.92 ± 0.81 | 17.0 | <0.0001 | 5.49 ± 0.34 | 4.98 ± 0.40 | -9.2 | 0.0013 |
| Striatum | 7.54 ± 1.51 | 8.18 ± 1.22 | 8.5 | <0.0001 | 8.59 ± 0.83 | 13.9 | <0.0001 | 5.36 ± 0.37 | 5.00 ± 0.50 | -6.7 | 0.0633 |
| Thal | 7.20 ± 1.34 | 7.90 ± 1.20 | 9.7 | <0.0001 | 8.33 ± 0.90 | 15.7 | <0.0001 | 6.68 ± 0.40 | 6.19 ± 0.43 | -7.4 | 0.0019 |
| VTA | 5.84 ± 1.13 | 6.38 ± 0.91 | 9.4 | <0.0001 | 6.66 ± 0.60 | 14.2 | <0.0001 | 5.14 ± 0.31 | 4.72 ± 0.50 | -8.2 | 0.0144 |
| VC | 6.93 ± 1.39 | 7.56 ± 1.25 | 9.1 | <0.0001 | 7.90 ± 0.95 | 14.1 | <0.0001 | 5.06 ± 0.43 | 4.62 ± 0.53 | -8.7 | 0.0085 |

NAc = Nucleus Accumbens; Amyg. = Amygdala; ACC = Anterior Cingulate Cortex; EntoC= Entorhinal Cortex; HIP = Hippocampus; AD = Anterior Dorsal; post = Posterior; Hypotha. = Hypothalamus; mPFC = medial prefrontal cortex; Olfact = olfactory bulb; OFC = Orbitofrontal Cortex; SSC = Somatosensory Cortex; Thal= Thalamus; VTA = Ventral Tegmental Area; VC = Visual Cortex.

**Figure 1.**
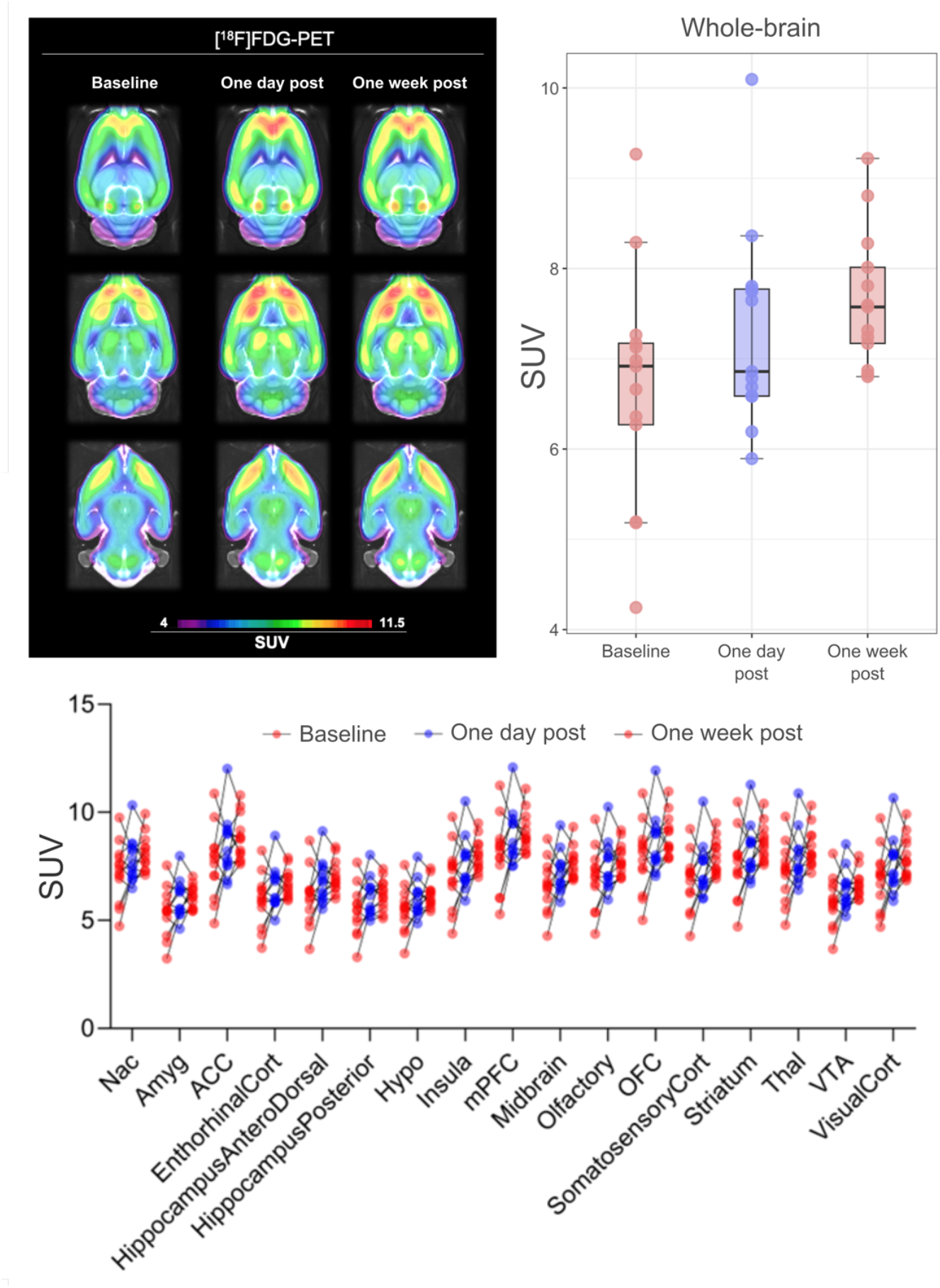
Change in global and regional [^18^F]FDG SUV following a single I.P. dose of rapamycin (8mg/kg) in 13 healthy Long-Evans rats. There was a significant increase in uptake in multiple brain regions following administration of rapamycin, which remained one-week after injection.

### [^18^F]SynVesT-1

For [^18^F]SynVesT-1 (n = 6), there was a significant decrease in pseudo-VT in multiple regions one day after rapamycin injection (Figure 2), which was also the case for a volume-weighted composite mask of all regions (p = 0.031, -7.8%). Regional decreases in binding ranged from -3.4 to -9.2%, with the strongest changes observed in the visual cortex and somatosensory cortex (Table 1).

**Figure 2.**
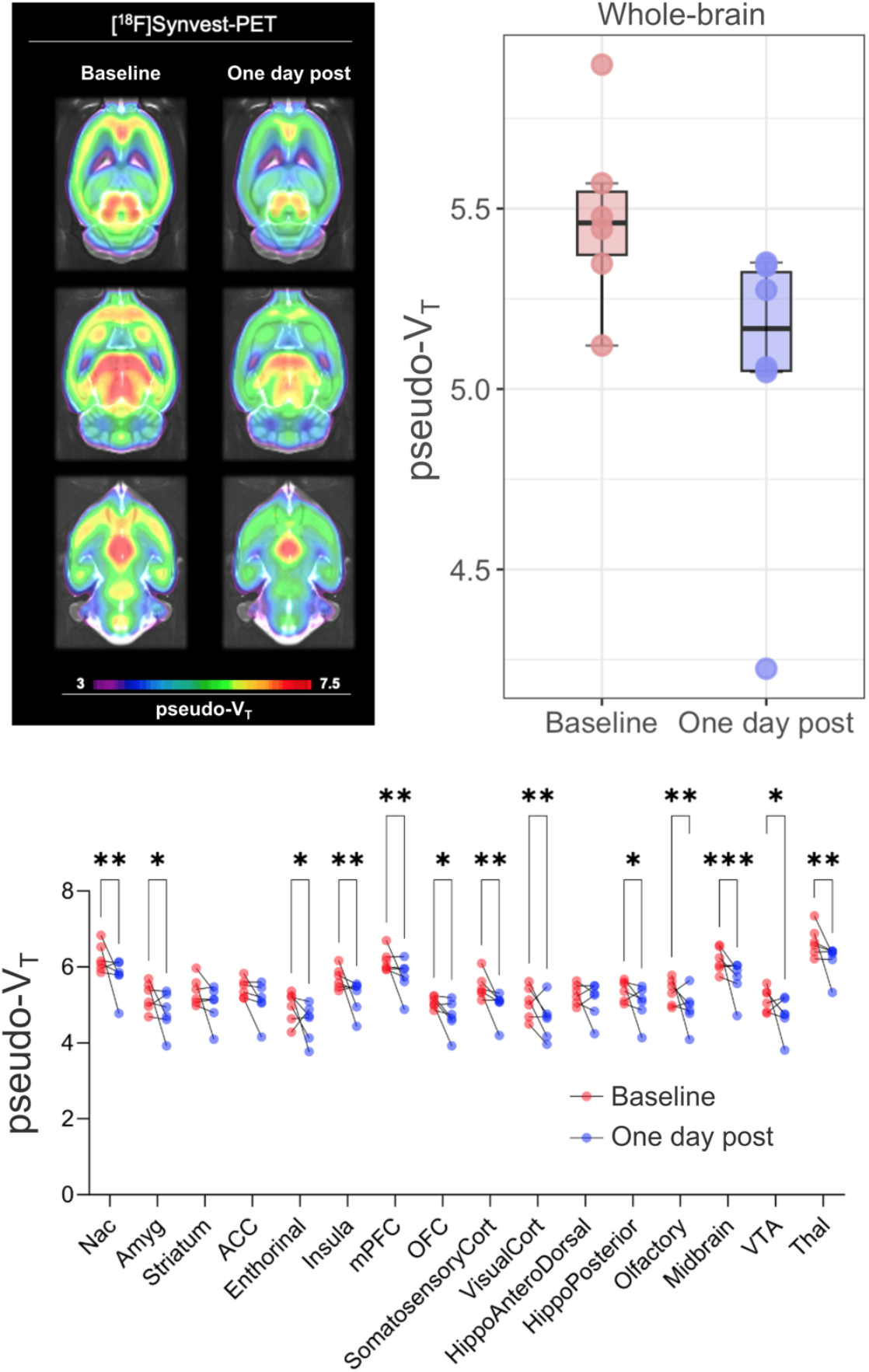
Change in global and regional [^18^F]SynVesT-1 pseudo-VT estimates following a single I.P. dose of rapamycin (8mg/kg) in 6 healthy Long-Evan rats. There was a significant decrease in binding in multiple brain regions the day after administration of the drug.

## DISCUSSION

In this study, we observed two distinct effects of single-dose rapamycin on brain function in young-adult Long-Evans rats: an increase in [^18^]FDG SUV at one-day follow-up that persisted for at least one week, and an acute decrease in [^18^F]SynVesT-1 binding at 24 hours post-administration. The sustained increase in glucose metabolism, as measured by [^18^F]FDG uptake, was evident across most brain regions examined, suggesting a broad effect of rapamycin on cerebral glucose metabolism. The reduction in [^18^F]SynVesT-1 binding, albeit observed in a small sample (n=6), indicates decreased synaptic density in multiple parts of the brain, though we did not collect longer-term data in this cohort.

The opposite changes we observed in metabolism and synaptic density indicate that rapamycin has divergent effects on brain processes. While our study design using separate cohorts for [^18^F]FDG and [^18^F]SynVesT-1 imaging precludes direct correlation between these measures, the findings point to potentially independent effects on energy metabolism and synaptic function. The increase in glucose metabolism may reflect enhanced cellular energy efficiency through mTOR inhibition, consistent with a previous study showing improved mitochondrial function and glycolysis following rapamycin treatment in AD mouse models [24]. The decrease in synaptic density could reflect an acute response to mTOR inhibition, as this pathway plays a key role in forming and regulating synaptic plasticity and protein synthesis [25,26]. These changes align with rapamycin’s established role in promoting autophagy and cellular stress resistance [27], processes that may temporarily reduce synaptic density while enhancing metabolic function.

Previous studies have demonstrated that mTOR signalling is essential for activity-dependent synapse formation in young neurons [28]. Importantly, the synaptic response to rapamycin appears to be age-dependent - while mTOR inhibition may suppress synaptogenesis and plasticity in young subjects, it could potentially have protective effects in aging brains where dysregulated mTOR signaling contributes to synaptic dysfunction and loss [29–31]. These differential age-dependent effects should be considered when interpreting our findings and evaluating the translational therapeutic potential of rapamycin in aged populations. However, we cannot exclude the possibility that rapamycin has detrimental effects on synaptic function, regardless of age. Further longitudinal *in vivo* imaging studies in older animals are therefore warranted.

These findings have implications for the development and assessment of rapamycin and other mTOR inhibitors as therapeutic compounds in neurodegenerative disorders. The lasting increase in glucose uptake is noteworthy since reduced cerebral glucose metabolism is a hallmark of AD and other neurodegenerative conditions [9]. Several clinical trials have been conducted, and are also being initiated, with the aim of evaluating rapamycin’s effects in patients with neurological disorders, such as multiple system atrophy [32], amyotrophic lateral sclerosis, [33] and early-stage AD [20]. The data from our pre-clinical experiments suggest that [^18^F]FDG and PET radioligands tracking synaptic density, such as [^18^F]SynVesT-1, could serve as response biomarkers in such trials, potentially providing early indicators of pharmacodynamic and therapeutic effects. The divergent effects we observed on metabolism and synaptic density also highlight the usefulness of complementary biomarkers to better characterize downstream effects of mTOR inhibitors.

Several limitations of our study should be considered when interpreting the findings. Importantly, both the [^18^F]FDG and [^18^F]SynVesT-1 experiments used a single-arm design without a placebo control group. While all animals received saline I.P. injections prior to scans where no rapamycin was administered, it cannot be excluded that the observed effects are caused to other factors than the drug itself, such as repeated anaesthesia. The single-dose design, while informative about acute effects, does not address the impact of a sustained rapamycin treatment that would be used clinically. The small sample size, particularly in the [^18^F]SynVesT-1 cohort (n=6), limits our ability to draw strong conclusions about subtle effects or regional variations in response. We also did not assess cognition, muscle or motor function in this study, and this lack of behavioral data prevents us from linking the observed changes in imaging estimates to functional outcomes. Additionally, the short follow-up period for synaptic density imaging leaves open questions about the observed changes beyond 24 hours. Future studies should address these limitations through longer-term treatment protocols, larger cohorts of older individuals, and comprehensive behavioral assessment. Understanding the relationship between changes in imaging and functional outcomes will be important for clinical translation, as will determining optimal dosing regimens that can maintain beneficial metabolic effects while avoiding potential negative impacts on synaptic function.

## ACKNOWLEDGEMENT

The study was supported by a Longevity Impetus grant from the Norn Group, Åke Wibergs Stiftelse (M24-0117 and M25-0306), Swedish Brain Foundation (PD2024-0444), the Karolinska Institutet Research Grants, the Lundbeck Foundation (R389-2021-1596), and the Neuroscience Academy Denmark.

## AUTHOR’S CONTRIBUTIONS

MP, PPS and JES conceived of the study and the design and provided the funding. MP, ZL and CB collected the data. FG, ZL and PPS performed the data analysis. PPS and MP supervised the study. PPS and FG drafted the manuscript. All authors revised the manuscript and approved the final version for publication.

## Notes

### Competing Interest Statement

The authors have declared no competing interest.

### Summary of Updates

Updated title, updated Fig 2., updated acknowledgement.

